# Using reference-free compressed data structures to analyse sequencing reads from thousands of human genomes

**DOI:** 10.1101/060186

**Authors:** Dirk D. Dolle, Zhicheng Liu, Matthew Cotten, Jared T. Simpson, Zamin Iqbal, Richard Durbin, Shane A. McCarthy, Thomas M. Keane

## Abstract

We are rapidly approaching the point where we have sequenced millions of human genomes. There is a pressing need for new data structures to store raw sequencing data and efficient algorithms for population scale analysis. Current reference based data formats do not fully exploit the redundancy in population sequencing nor take advantage of shared genetic variation. In recent years, the Burrows-Wheeler transform (BWT) and FM-index have been widely employed as a full text searchable index for read alignment and de novo assembly. We introduce the concept of a population BWT and use it to store and index the sequencing reads of 2,705 samples from the 1000 Genomes Project. A key feature is that as more genomes are added, identical read sequences are increasingly observed and compression becomes more efficient. We assess the support in the 1000 Genomes read data for every base position of two human reference assembly versions, identifying that 3.2 Mbp with population support was lost in the transition from GRCh37 with 13.7 Mbp added to GRCh38. We show that the vast majority of variant alleles can be uniquely described by overlapping 31-mers and show how rapid and accurate SNP and indel genotyping can be carried out across the genomes in the population BWT. We use the population BWT to carry out non-reference queries to search for the presence of all known viral genomes, and discover human T-lymphotropic virus 1 integrations in six samples in a recognised epidemiological distribution.

## INTRODUCTION

Recent years have seen the number of whole human genomes sequenced continue to increase dramatically through large scale population and medical sequencing projects such as the 1000 Genomes Project (The 1000 Genomes Project Consortium 2015), UK10K (The UK10K Consortium 2015), and GoNL (The Genome of the Netherlands Consortium 2014). The scale up of human population sequencing has enabled us to detect sequence variants down to extremely low minor allele frequencies (The 1000 Genomes Project Consortium 2015), explore variation in ancient human lineages and isolated populations (Raghavan et al. 2015), and use genomics to discover rare disease causing mutations(Katsanis and Katsanis 2013). Current predictions estimate that we will have sequenced 1M human genomes in the near future (Stephens et al. 2015), which will present formidable informatics scaling challenges.

The sequencing data produced by current high throughput sequencing technologies consists of paired reads on the order of a hundred base pairs along with their base qualities with the vast majority of aligned data currently stored in the SAM/BAM format (Li et al. 2009). The SAM/BAM format, originally developed by the 1000 Genomes Project (1000GP), requires on the order of one byte per base pair with the vast majority of the space being taken by the base qualities (Hsi-Yang Fritz et al. 2011). Recently, the CRAM format has been proposed (Hsi-Yang Fritz et al. 2011) and adopted by the Global Alliance for Genomics and Health consortium (https://genomicsandhealth.org/) to provide a more sustainable foundation for exploring strategies for controlled loss of base qualities, a strategy that can result in more accurate genotyping (Ochoa et al. 2016; Yu et al. 2015). One key innovation of the CRAM format is to only store the differences in individual sequencing reads relative to the reference genome. Furthermore, when one considers that the vast majority of variants per individual are shared amongst multiple individuals (The 1000 Genomes Project Consortium 2015), there is also significant duplication of non-reference sequences.

In parallel, there is increasing interest in methods for rapid searching of large collections of sequencing reads from many individuals. Iqbal *et al.* (2012) developed the Cortex assembler for representing sequencing reads from multiple samples using coloured de Bruijn graphs for genome assembly and reference-free variant identification (Iqbal et al. 2012). Applications that were presented include variant calling from a single high-coverage genome, detection of novel sequence from a population not present in the reference, and genotyping of simple and complex variants highly divergent from the reference. However, the implementation only scaled to around 10 human genomes on standard hardware, orders of magnitude lower than what is required. Recently, Bloom filters in the form of Sequence Bloom Trees (SBTs) were used to build highly compressed partial text indexes given a large set of input sequences and demonstrate rapid sequence searches with low memory requirements (Bloom 1970; Solomon and Kingsford 2016). An SBT structure was constructed from 2,652 RNA-seq experiments, requiring 200 GB. In recent years, the Burrows-Wheeler Transform (BWT) and FM-index have been widely employed to build full-text indexes for read alignment (Langmead et al. 2009; Li and Durbin 2009), read error correction (Li 2015), and *de novo* genome assembly (Simpson and Durbin 2011). The key features of using a BWT structure to index sequencing reads is that it is inherently reference-free, full-text, compressed, and coupled with the FM-index enables rapid sequence searches of arbitrary k-mer sizes across the entire set of sequences without rebuilding the index for different values of k.

In this paper, we explore the use of the BWT structure and FM-index to build a full-text index of the sequencing reads from 2,705 individuals across 26 populations from the 1000GP (The 1000 Genomes Project Consortium 2015). We show that there exists significant duplication of sequence across the populations and how the data can be reduced to a small fraction of unique sequences. A key property of this strategy is that as more whole human genomes are added, the growth rate of the total size of the structure decreases, since each new genome only adds a small fraction of new unique sequences. We use the structure to carry out a single base resolution comparative analysis of two recent versions of the human reference assembly, using the BWT to determine the population support at every position. We also show how the population BWT can be used to carry out rapid and accurate SNP and indel genotyping across the entire population, and how the reference free nature of the structure enables rapid searching for non-reference sequences such as viral sequences.

## Results

### Data processing and BWT construction

Figure 1 gives an overview of the data processing strategy. We begin with the whole-genome low coverage and exome sequencing reads from the final phase of the 1000GP (2,705 individuals over 26 populations) consisting of approximately 87 Tbp and 922 billion reads (The 1000 Genomes Project Consortium 2015). We used a combination of examining the base qualities for each read and querying the sequences against a pre-constructed Cortex graph (Iqbal et al. 2012) to carry out error correction and removal of poor quality reads (see Methods). This resulted in a set of 734 billion unchanged reads, 85 billion corrected reads, and 103 billion reads that could not be corrected and were discarded. We took advantage of the reference strand labelling in the Cortex de Bruijn graph (obtained by labelling nodes during a traversal of the reference sequence) to reverse complement read sequences with a clear reverse strand orientation with respect to the reference genome (see methods). We normalised the read lengths by trimming to 73 or 100 bp depending on whether the original read sequence was greater than 100 bp. For each resulting read, we used a key-value pair database (RocksDB, http://rocksdb.org/) to record the read groups, number of corrected bases, and number of bases greater than Q20 using the read sequence as the key. We next sorted the 53 billion sequence keys reverse lexicographically and constructed the BWT structure. The 53 billion unique sequences (keys) produced an average of 15.45 reads for each key. In Figure 1d, we benchmarked the total size of the BWT using both the uncorrected and corrected reads with increasing numbers of individuals (using the reads from a 5Mbp region on chr20). The plot shows that using the uncorrected reads, the BWT continues to linearly increase in size, independent of the sort order. Reverse lexicographic sorting order (RLO) performed an order of magnitude better than lexicographic order (LO). The effect of error correction of the reads can be observed with the total BWT around two orders of magnitude larger with uncorrected reads. With error correction and RLO sorting, the total size of the BWT begins to plateau from approximately 1,500-2,000 genomes. The final size of the BWT for the entire dataset was 464 GB (split over 16 smaller BWTs based on read prefix in order to load into system memory over multiple servers, Supplementary Table 1) and the corresponding RocksDB metadata database was 4.75 TB (0.09 bytes per bp). The resulting population BWT server can be queried for exact matches to arbitrary length k-mer sequences and return either the count of matching read sequences, the matching read sequences, or the matching read sequences with sample metadata (Supplementary Figure 1). We benchmarked the query completion time for 100,000 k-mer queries for the different types of server responses and k values (Supplementary Table 2), finding that for smaller values of k returning matching read counts was the fastest primarily due to the network time required for transferring large quantities of read sequences. At larger values of k where less matching reads are found, the difference between requesting read counts and full matching read sequences is considerably reduced. In the remainder of the paper, we call the resulting population BWT the 1000GP BWT or where unambiguous just the BWT.

**Figure 1.**
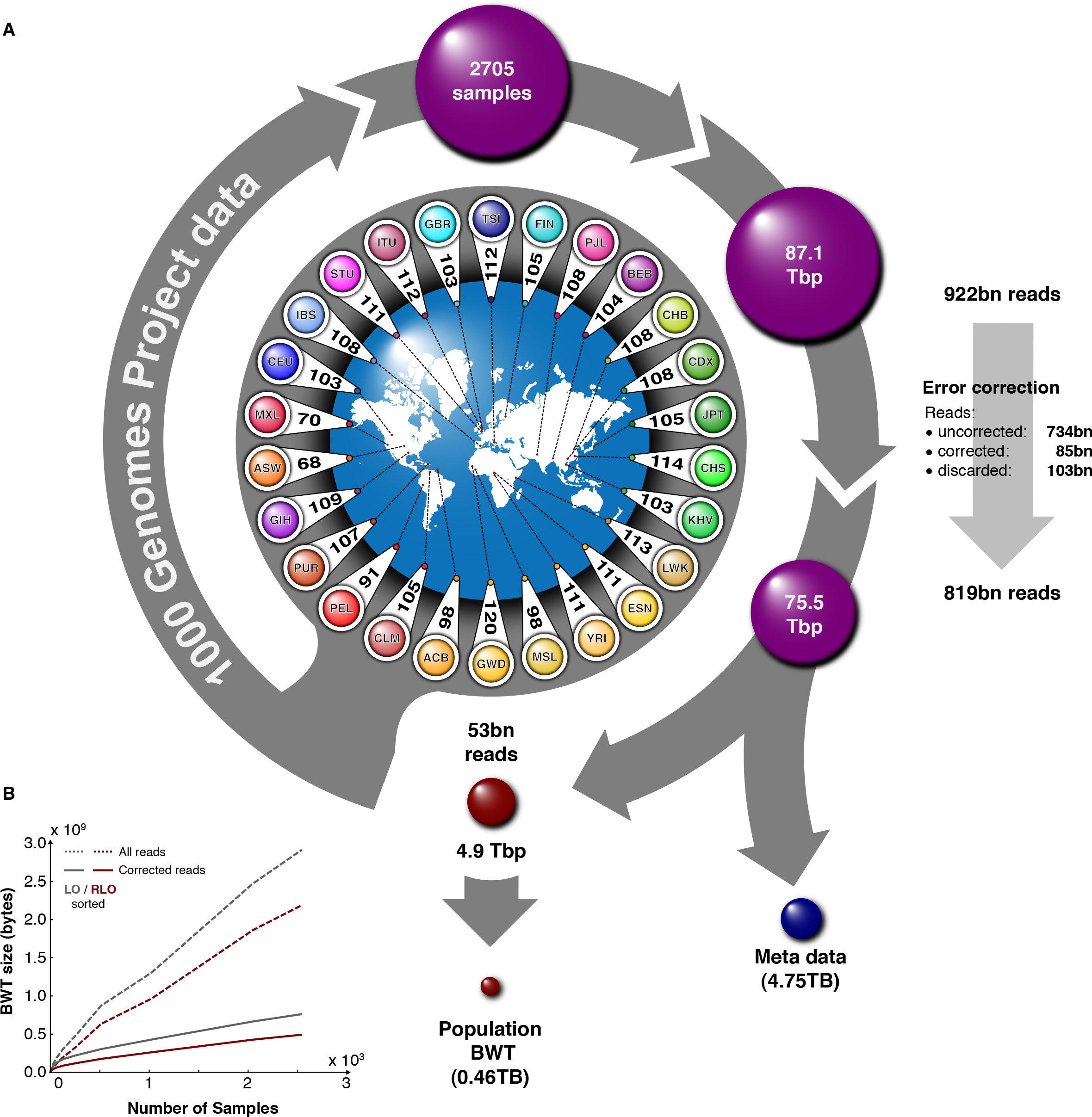
(a) Sequencing reads from 2,705 individuals (low coverage whole-genome and
exome sequencing) from 26 populations comprising a total of 922 billion reads (87.1 Tbp) used for the 1000GP population BWT. Reads were first error corrected using a Cortex graph (Iqbal et al. 2012). The error corrected reads were then trimmed to either 100 bp or 73 bp, unique sequences identified on the forward strand, quality values discarded and the metadata stored in a separate database. This resulted in 4.9 Tbp consisting of 53 billion non-redundant reads. (b)Sequences were sorted by reverse lexicographic order to build the population BWT. Different sorting orders were tested for their effect on the BWT size using the 1000GP reads aligned to a 5 Mbp region.

### Population support for human reference assemblies and variation

We first used the population BWT to assess the direct support in the 1000GP read data for every base of two recent versions of the human reference assembly (GRCh37 and GRCh38) and support for the SNP and short indel variants called by the 1000GP. We extracted all forward strand 31-mers contained in both reference assemblies and queried the population BWT for reads matching these 31-mers. Finally, we also generated all 31-mers contained in the reads stored in the BWT Read Server. The vast majority of reference 31-mers (Figure 2a) are supported by the 1000GP BWT (99.97%) and mostly shared between both assemblies (99.41%) with 0.07% of GRCh37 31-mers lost from the change from GRCh37 to GRCh38 with 0.49% gained in GRCh38. We further queried the 1000GP BWT for all 31-mers generated by the SNP and indel variants found by the 1000GP (The 1000 Genomes Project Consortium 2015). Figure 2b shows the intersections of these four 31-mer sets. Considering the reference genomes and the 1000GP variant 31-mers, the vast majority of 31-mers were either reference (solid black outline) or variant specific (dotted outline) and supported by the 1000GP BWT (overlap with the red ellipse). Figure 2c shows the amount of sequence gained or lost over four functional categories based on the GENCODE human genome annotation (Harrow et al. 2012). When the 31-mers are converted into reference genome regions, 3.1Mbp (1.6M 31-mers, 0.07%) of sequence that has population BWT support was lost in the transition from GRCh37 to GRCh38 but roughly 7.5 times (13.6Mbp) more was gained (10.5M 31-mers, 0.49%). We examined the read coverage for the regions in GRCh37 that do not contain 31-mer support from the 1000GP BWT. The vast majority of these regions are 50-60bp (Supplementary Figure 2), with more than 70% (89% and 73.7% for GRCh37 exclusive regions and those shared with GRCh38, respectively) overlapping at least one variant. 65% and 39% (for GRCh37 exclusive regions and those shared with GRCh38, respectively) overlap a locus for which GRCh37 contains the minor allele or an error. Interestingly, the majority of the unsupported GRCh38 sequence is located in the new synthetic centromeric regions (CTM, Figure 2c: 242 Kbp) although 2.8 Mbp of the new centromere is supported. The amount of coding sequence without population support in GRCh37 consists of 9.2 kbp in 203 protein coding genes and 4.6 Kbp in 123 genes for GRCh38 (Supplementary Table 4), reflecting the flipping of bases to the major allele. Interestingly, there were 12 protein coding genes that contain unsupported 31-mers only found in GRCh38.

**Figure 2.**
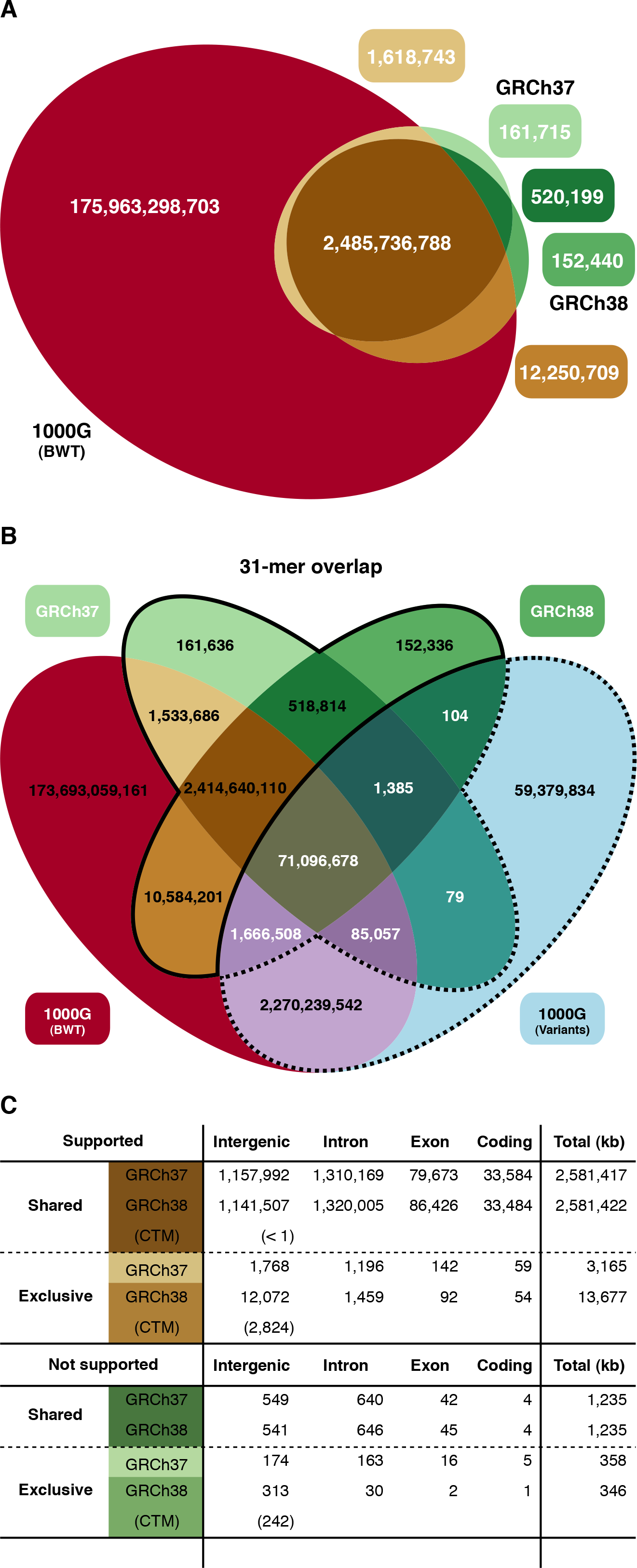
(a) 31-mer intersection of two human reference assemblies (GRCh37 and GRCh38) and the 1000GP population BWT (b) 31-mer intersection of two human reference assemblies, 1000GP population BWT, and all 31-mers generated from the 1000GP phase 3 SNP and indel variants (The 1000 Genomes Project Consortium 2015). 31-mers shared between reference sets and variant set (white numbers) make up for approximately 3% of each data set and almost all (99.998%) are supported by the 1000GP population BWT. (c) A breakdown of the regions on the two human assemblies with and without 1000GP population BWT support that is shared or exclusive to either genome build (all numbers are kbp), in four functional categories. CTM refers to the centromeric sequence.

### Reference-free population genotyping

Figure 2b shows that the majority (97%) of 31-mers derived from 1000GP variation catalogue are distinct from the reference genome. Furthermore, we determined that 99% of these 1000GP variant 31-mers not found in the references are locus specific (no other combination of variants in either the same or a different locus in the GRCh37 assembly generates the identical 31-mer), Figure 3a. The 31-mers shared between the reference genomes and the variants are likely to be in regions containing repeats longer than 31bp, which were still callable in the 1000GP by using the untrimmed longer reads or read pair information. Informed by this analysis, we developed a simple SNP and indel genotyping strategy based on querying the population BWT for k-mer sequences to test for read support of the reference and alternative allele for every individual. We tile across each genotyping site with overlapping k-mers upstream and downstream of the site, and query the population BWT for exact matching reads. We assign a genotype to each sample by recording how many of the reads from the sample match best to the reference or alternative allele (see Methods).

**Figure 3.**
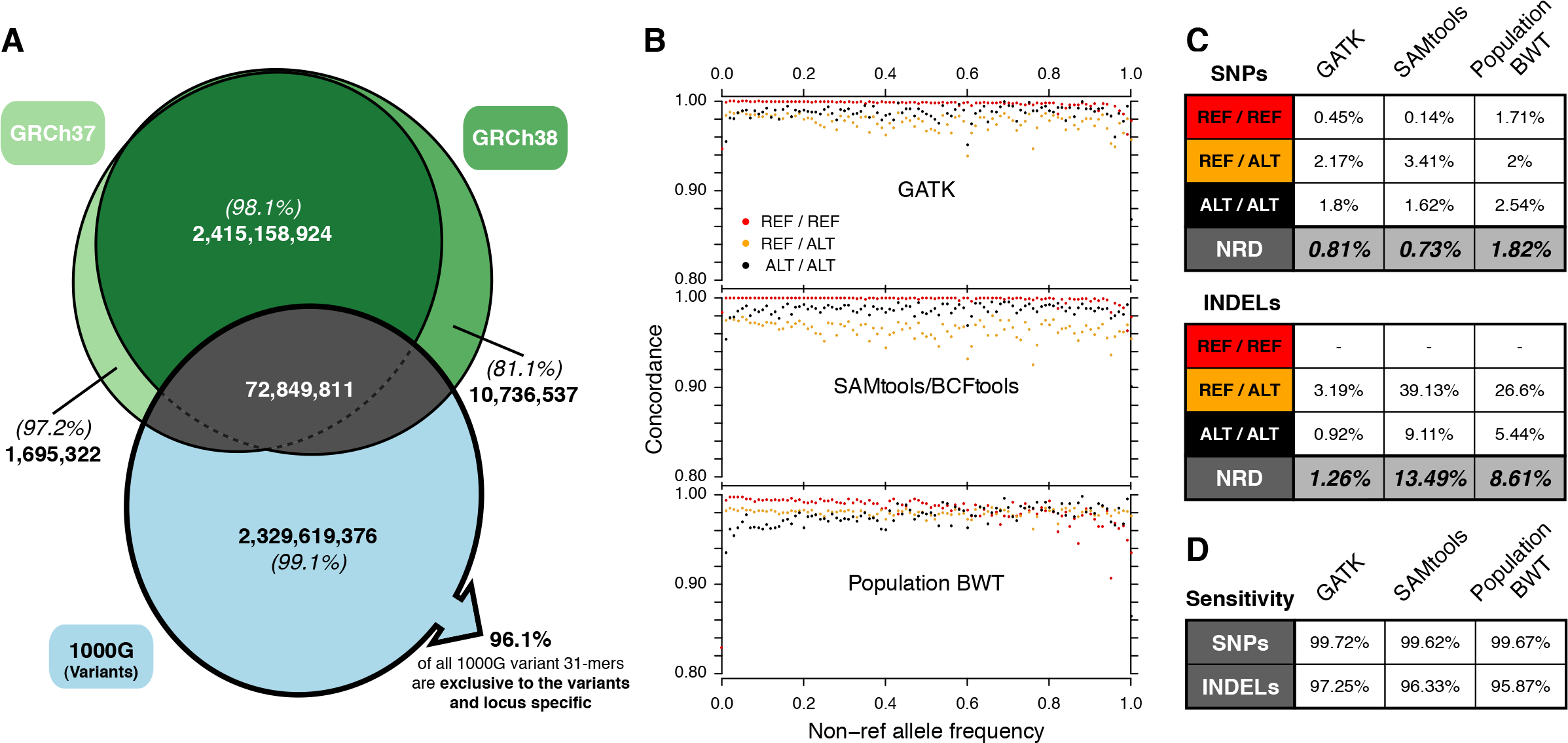
(a) Intersection of the human reference assembly 31-mers and the 1000**GP SNP** and indel variant 31-mers. The percentages in brackets give the proportion of these 31-mers that are locus specific (no other combination of variants in either the same or a different locus in the GRCh37 assembly generates the identical 31-mer). 96.1% of all 31-mers generated based on 1000GP variants are locus specific and exclusive to the variants set with 91.8% containing a single alternative allele. (b) SNP genotyping of the 1000GP samples at Illumina Omni chip exome-only sites by 31-mer querying of the BWT compared to single sample calling with GATK HaplotypeCaller (v3.5) and Samtools (v1.1). Dots indicate genotype concordance for variants at different allele frequencies. (c) Genotype discordance rates for SNPs (Omni exome-only: 80,973 sites, all samples) and indels (Genome in a Bottle (Zook et al. 2016) exome in NA12878: 654 sites). (d) Sensitivity of each method expressed as the fraction of total genotypes for which a genotype call was made.

For SNPs, we benchmarked the approach using the Illumina Infinium BeadChip Omni2.5-8 genotypes in the 1000GP exome regions as a truth set. We initially evaluated the effect different values of k have on the population BWT genotyping accuracy using all chromosome 20 sites, finding that k=34 produced the lowest non-reference discordance (Supplementary Table 3). We genotyped all of the Omni chip positions in the 1000GP exome regions with single sample calling using GATK HaplotypeCaller, Samtools/BCFtools, and the 1000GP population BWT. Figure 3b shows that the population BWT genotyping compares favourably to GATK and Samtools across the allele frequency spectrum. The overall non-reference discordance rate is slightly higher for the population BWT genotyping (1.82%) compared to the GATK (0.81%) and Samtools (0.73%). For heterozygous SNPs, the population BWT approach is more accurate than the two reference based callers (discordance rate of 2% vs. 2.17% for GATK, and 3.41% for Samtools). The proportion of sites genotyped was greater than 99% for all three approaches, Figure 3d.

We developed a similar approach for indel genotyping by testing reference and alternative alleles by dense k-mer tiling across the indel site (see methods), querying the population BWT with the resulting k-mers, and assigning a genotype to each individual based on the matching reads returned (see methods). For indels, we use the Genome in a Bottle (GIAB) consortium gold standard indel genotypes for NA12878 for evaluation (Zook et al. 2016). Initially, we tested the effect different values of k have on genotyping accuracy using chromosome 20 sites, determining that k=25 produced the most accurate genotypes (Supplementary Table 3). We genotyped the indels with GATK HaplotypeCaller, Samtools/BCFtools, and the 1000GP population BWT (see methods). The indel genotyping accuracy varied widely between the callers. GATK produced the lowest non-reference discordance (1.26%), followed by the 1000GP population BWT (8.61%), and Samtools (13.49%), Figure 3c. This is not so surprising since the GIAB indel calls are largely derived from GATK genotypes and there is often poor overlap between indel discovery tools (Narzisi et al. 2014).

### Non-reference queries

As the population BWT is a full text index of the read sequences, irrespective of whether they align to the reference genome or not, it enables rapid testing of hypothesis driven queries. We sought to assess the proportion of sequences of viral origin contained in the 1000GP reads. An earlier study using 150 individuals from the 1000GP, found evidence for 0.13% of reads coming from non-human DNA (Tae et al. 2014). To expand this to the full set of samples, we downloaded 257,943 viral sequences from the CoreNucleotide division of GenBank and used the Kraken classifier (Wood and Salzberg 2014) to define a set of 102.6M virus specific 31-mers (Figure 4a) (see methods). We queried the 1000GP population BWT with these 31-mers initially for read counts (to remove very highly abundant low complexity sequences), then returned matching read sequences, and finally queried the metadata database for sample information. The population BWT queries were run in under two days, with the sample metadata retrieval taking seven days (see methods). The most prevalent source of non-human sequences is the Herpesviruses, including Epstein-Barr virus, used in the creation of the Lymphoblastoid cell lines (LCLs) that were the DNA source for many of the 1000GP samples. The distribution of the number of EBV matching reads largely follows the documented DNA source in the 1000GP (Figure 4b), with a few notable exceptions which are likely misclassified as being from blood. The DNA that is recorded as being of unknown origin appear to be almost entirely from LCLs, having a similar distribution of EBV reads as the documented LCL derived samples. Of the viruses identified (excluding EBV), 69 occur in at least one sample at greater than 10 reads, and 14 at greater than 100 reads (Supplementary Figures 2-7).

**Figure 4.**
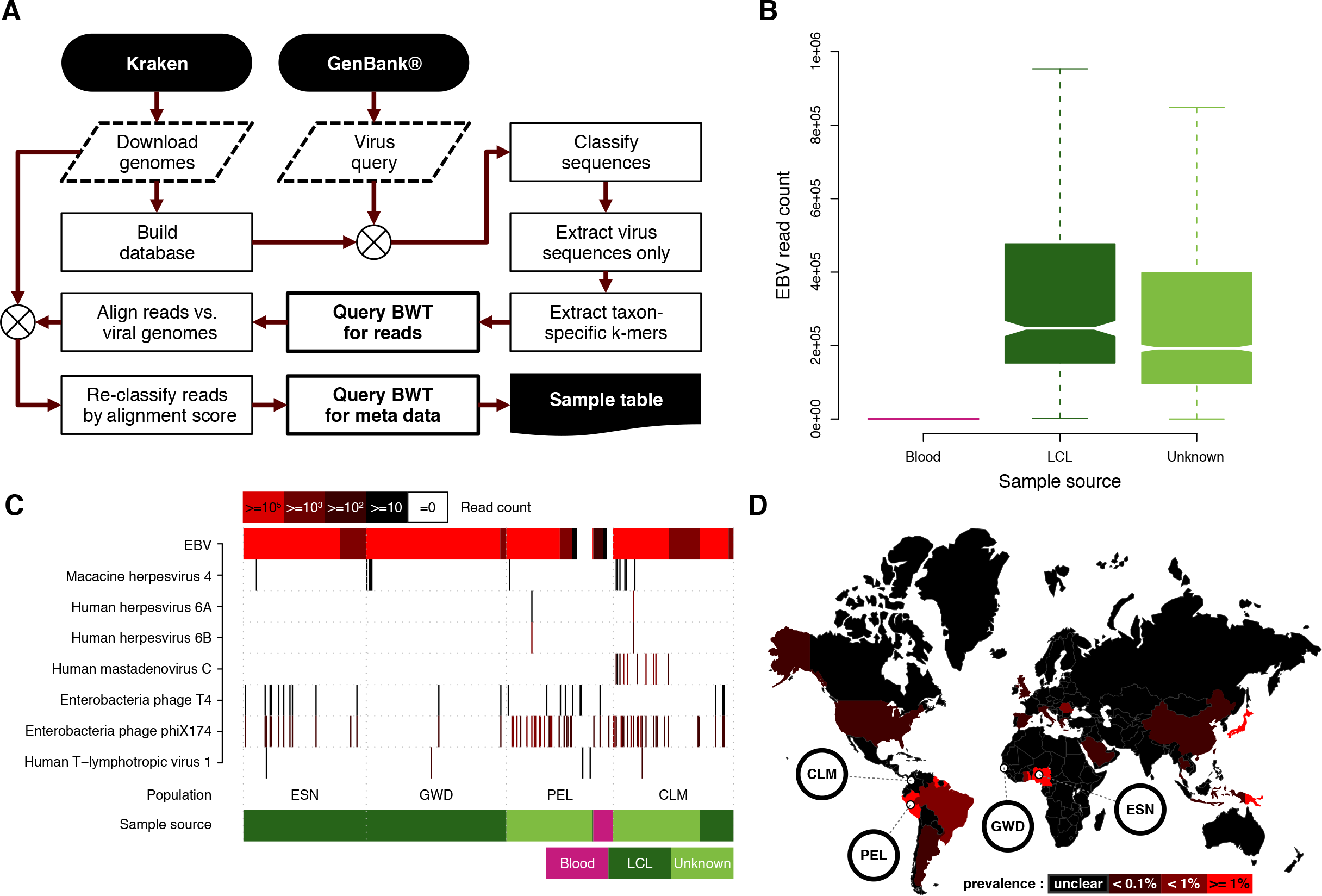
(a) Reference genomes (Human, bacteria, plasmids, and viruses) were downloaded using Kraken’s (Wood and Salzberg 2014) built-in routines and a Kraken database generated.GenBank was queried for all virus sequences and the resulting sequence set classified using Kraken to identify taxon specific 31-mers which were used to query the population BWT for matching reads. Retrieved read sequences were re-classified by alignment to the viral genomes stored in the Kraken database. Finally, sample metadata was retrieved for the final read set. (b) Notched boxplot showing the distribution of human herpesviruses (including EBV) read counts stratified by documented DNA source. Not-overlapping notches indicate a significant difference of the medians at the 5% level. (c) The populations for which at least one sample contains greater than 10 HTLV-1 reads (black bars) and other virus taxa with greater than 99 reads (red bars) in at least one sample are shown (for all populations, see Supplementary Figs. 3-8) (d)World map showing HTLV-1 prevalence in different countries (adapted from (Verdonck et al.2007)) with 1000GP populations that show signal for this virus highlighted.

Figure 4c gives the species source for the most frequently found sequences per individual for four particular population groups (see Supplementary Figs. 3-8 for all populations).Enterobacteria phage phiX174, the Illumina library spike-in sequence, is also prevalent across all of the populations at greater or equals 100 reads in 605 samples. Adenovirus C is reported to be present in almost all populations worldwide (Garnett et al. 2002), however our analysis shows that it is almost completely absent several populations (e.g. Gambian in Western Divisions in the Gambia, and Esan in Nigeria, in Figure 4). The absence of adenovirus in some groups and high levels in other groups suggests differences in the sample preparation (Adenovirus C is often used as a recombinant vector for cell culture reagents (Luo et al. 2007) or differences in adenovirus in these populations.

One interesting finding is the presence of Human T-lymphotropic virus 1 (HTLV-1) reads found in six individuals (Figure 4c and (Table 2). HLTV-1 can integrate into the genome and is known to have infected human populations for thousands of years with the virus being transferred from mother to child, through sexual contact, or contaminated blood products (Derse et al. 2007; Verdonck et al. 2007). The known epidemiological distribution spans areas of southern Japan (Satake, Yamaguchi, and Tadokoro 2012), sub-Saharan Africa, the Caribbean, and South America where more than 1% of the general population is infected (Verdonck et al. 2007). For the most part, carriers remain asymptomatic but HTLV-1 infection has been associated with exceptionally severe diseases, such as adult T-cell leukaemia/lymphoma (Takatsuki 2005) and an inflammatory disease of the central nervous system called HTLV-1-associated myelopathy/tropical spastic paraparesis (Gessain et al. 1985; Lezin et al. 2005). The 31-mer querying method identified six samples from five populations with potential HTLV-1 integrations. For those individuals, we also aligned the entire original read set to a reference genome containing GRCh38 and a HTLV-1 consensus sequence, confirming the presence of HTLV-1 in these genomes and slightly increasing the HTLV-1 read support for each sample (Table 2). The populations in which we detected HTLV-1 presence largely follow the known epidemiological distribution with HTLV-1 positive samples from Africa and South American populations, and were sequenced at six different centres. We did not observe HTLV-1 in any Japanese samples (reported HTLV-1 prevalence of 0.66% and 1.02% (Satake et al. 2012)) although Japan has had HTLV-1 population screening in place since 1986 (Inaba et al. 1989).

## Discussion

In this paper, we show how BWT indexes can be used for efficient compression and indexing of large collections of sequencing reads from thousands of individuals. Unlike traditional reference based alignment approaches, the population BWT has a sub-linear growth as more individuals are included in the structure. However, this is dependant on the sequencing data having a low error rate so that the majority of new sequences observed in each individual represent true genetic variation. One of the main difficulties of using the 1000GP data with this approach is that most individuals were sequenced to low coverage (7-8x). For error correction, we used a Cortex de Bruijn graph that was built from these reads and was error-cleaned by removing tips (short contigs unconnected at one end) and unitigs which were at low frequency in all populations. The fraction of error corrected reads was quite low (9.2%) since our error correction strategy was deliberately conservative as we wanted to avoid removing true genetic variation. It is still notable that the resulting population BWT contains over thirty-five times more 31-mers than are present in the reference genomes and the SNP and indel variants (Figure 2a). It has been suggested that existing variation catalogs fail to account for 35-68% of some types of structural variation and 25% of short indels (Gordon et al. 2016). Therefore, unaccounted genetic variants, variants located in inaccessible regions of the genome, and non-human sequences could contribute to these novel 31-mers. Our virus sequence analysis only accounted for 102M 31-mers in the population BWT, therefore it is more likely that these novel 31-mers are due to remaining errors in the sequencing reads. One could perform more stringent error correction to reduce the sequencing errors at a cost of removing true low frequency variants. More recent approaches to per-sample read error correction are most effective with comparatively high sequencing depth (30-50x) per sample (Li 2015; Simpson and Durbin 2011). Therefore, we envisage that as the cost of human sequencing continues to decrease and higher depth sequencing becomes the norm, the population BWT could be an efficient storage medium for indexing large collections of human samples.

The most significant storage saving in this approach comes from discarding the base qualities after base error correction is carried out. It remains an open question as to what proportion of base qualities need to be retained for accurate variant discovery and genotyping, with increasing evidence showing that discarding or quantile binning of base qualities does not have a detrimental effect (Ochoa et al. 2016; Yu et al. 2015). However, many applications of next-generation sequencing (e.g. clinical sequencing) rely on highly accurate identification of novel rare variants. One alternative approach to completely discarding base qualities could be a controlled loss of base qualities. For example, there could be an iterative process of population BWT construction where genomes are continually added. Initially with few genomes, the majority of the sequencing reads will contain novel k-mers and as more genomes are added, we will observe the same k-mers in multiple individuals across the population. One could envisage an approach where base qualities are only maintained for reads that support novel kmers with these k-mers being constantly queried against the BWT for increasing population read support with the goal of eventually discarding these base qualities as increasing support is observed in the population. One could employ the BEETL-fastq BWT based data structure to create a side structure of compressed and searchable indexes of read sequences including base qualities (Janin, Schulz-Trieglaff, and Cox 2014).

One of the limitations of this approach is that this implementation of a population BWT does not maintain read pair information. In our SNP and indel genotyping, read pair information could be incorporated into the genotyping strategy to derive more accurate genotypes. Read pairs would be particularly useful for structural variant discovery and genotyping as most existing structural variation detection algorithms use a combination of split reads and read pairs for supporting evidence (Keane et al. 2014; Layer et al. 2014). In our virus analysis, efficient retrieval of read pairs would enable more rapid localisation of the HTLV-1 viral integrations by avoiding the need to realign the full original read set. For SNP and indel discovery, retrieval of read pairs would enable local haplotype assembly and phasing of discovered variants which could aid correct alignment of highly variable loci into a variation graph (Church et al. 2015), especially with library technologies that conserve long range phase information (Putnam et al. 2016; Zheng et al. 2016). Recording read pair information could significantly increase the amount of metadata required from just basic sample level information to knowledge of every unique read pair combination or sets of reads from the same molecule. The ability to store and efficiently retrieve all of the read pairs of a sample could enable the use of the population BWT as a highly compressed, searchable, and scalable archival format for sequencing data.

One of the benefits of choosing the BWT and FM-index as the underlying data structure is that the construction process does not constrain the length of possible k-mer queries. In de Bruijn based approaches such as Cortex (Iqbal et al. 2012) and SBT (Solomon and Kingsford 2016), the k-mer must be fixed at the time of index construction. In the SNP and indel genotyping, the genotyping accuracy varied depending on the k-mer. The length of the k-mer used to assess an individual site can be affected by the number of mutations in the local region, where smaller, more densely sampled k-mers could potentially produce more accurate genotypes in regions of high mutation rates. Using dynamic k-mer queries for genotyping and the incorporation of read pair information are potential avenues for further improving genotyping accuracy.

Using whole-genome sequencing reads to classify reads into taxonomic groups has become the basis for metagenomic analysis (Gilbert and Dupont 2011). We used a metagenomics k-mer classification approach to detect evidence for non-human sequences in the 1000GP reads. Several studies that have cautioned against over interpretation of unexpected sequences found in sequencing reads due to possibility of laboratory kit or reagent contamination (Lusk 2014; Salter et al. 2014). For these reasons, our finding of evidence for low levels of HTLV-1 in several 1000GP samples should be treated with caution. On the one hand, the epidemiological distribution of the samples found to contain HTLV-1 fits the known pattern, we can localise many of the putative integrations using read pairs (Table 2), and the samples were sequenced at multiple different centres. However, we cannot rule out possibility of kit or reagent contamination without further laboratory validation of the results.

## Methods

### Sequencing data

The sequencing reads were downloaded in fastq format from the 1000GP ftp site and correspond to the phase 3 sequencing data freeze (ftp://ftp.1000genomes.ebi.ac.uk/vol1/ftp/phase3/20130502.phase3.analysis.sequence.index) consisting of 2,574 in total and 2,535 of these with both low coverage whole-genome and exome sequencing (The 1000 Genomes Project Consortium 2015).

### Error correction

Read error correction was carried out using the Cortex software (Iqbal et al. 2012) (https://github.com/iqbal-lab/cortex). Briefly, Cortex is a de novo De Bruijn graph assembler that allows simultaneous assembly of multiple samples and variants to be called without reliance on mapping of reads to a reference genome. We used the Cortex graph that contains a merge of all of the populations in the 1000GP (ftp://ftp.1000genomes.ebi.ac.uk/vol1/ftp/technical/working/20130718_phase3_samples_cortex_graphs/phase.all_pops.ctx) (see Supplementary Methods of (The 1000 Genomes Project Consortium 2015)). The Cortex graph was loaded into memory, and then the reference genome (GRC37) was parsed, annotating each kmer with the direction in which it was seen (forward, reverse or both). If a read was less than 73bp in length or contained any character other than ACGT, it was discarded. If all of the base qualities for a read were greater than or equal to Q20, the read was kept without correction. Correction was seeded by finding a 31-mer of Q>20 bases, and extending greedily by shifting one base at a time. On shifting and meeting a Q<20 base, if there was precisely one single-base correction of a Q<20 base which changed a kmer absent from the Cortex graph to a kmer present in the Cortex graph, this change was made. If all of the kmers in a read were annotated consistently with the read coming from the reverse strand of the reference genome (i.e. either unannotated, or annotated as being seen in the reverse strand of the reference), the corrected read was reverse-complemented and printed in the forward direction, otherwise it was printed in the same orientation as the input data. This was done purely to improve compression in the BWT. Finally, read sequences were trimmed to two reads lengths: 73 and 100bp. If a corrected read was greater than 100bp, then it was trimmed to 100bp; if a read was between 73bp and 100bp in length, it was trimmed to 73bp; and if a read was less than 73bp, it was discarded. For base error correction, we used a modified version of error_correction.c from Cortex (https://github.com/wtsi-svi/cortex@fc26874).

### Read deduplication and metadata

The error correction process output the read sequence (in forward orientation), the read name, the number of corrected bases during and the number of low quality (<Q20) bases. Corrected read sequences were sorted in Reverse Lexicographic Order (RLO) with duplicates removed. For each unique read sequence in the final read set, we stored the read groups (2 bytes), and the number of corrected bases (1 byte) and the number of low quality bases (1 byte). This information was stored in a RocksDB (v2.6) with the unique read sequences as keys.

### BWT and FM-index construction

The reads were split into 16 partitions based on the last two base pairs in the read sequence (see Supplementary Table 1) with the reads for each partition sorted in reverse lexicographic order. Then we used SGA v0.10.13 (Simpson and Durbin 2011) to construct the BWT string for each read collection. SGA outputs BWT strings in Run-Length Encoding (RLE) with each byte representing a continuous run of the same character. The first three bits of a byte encode the five different characters (i.e. ACGT$). The last five bits of the same byte encode the number of the runs for that character up to the length of 31. The cumulative size of the run-length encoded BWTs on disk was 464GB.

The Burrows-Wheeler Transform renders an important property Last-to-First (column) mapping, i.e. the ith occurrence of character X in the last column corresponds to the ith occurrence of X in the first column. The FM-index (Ferragina and Manzini 2000), based on BWT and LF mapping, allows for fast query of a pattern and locate every occurrence of the searched pattern. We built an index structure based on the Run-Length Encoded BWT string. With such index, we were able to search for a kmer, extend to the full read from matched location and get the full read sequence in linear time. The implementation of this index can be found at https://github.com/wtsi-svi/ReadServer.

### System setup

We set up a server to allow fast query of a given k-mer and return information about the number of matched reads, the matched read sequences, and for each matched read, the list of samples that the read was derived from. To achieve high throughput and fast response, we created a message queue based application server that sends k-mer sequence requests to the 16 BWTs across four physical servers (Supplementary Figure 1). Each machine has four applications and each application has a BWT partition and its associated index structure loaded in memory (total memory required: 561GB). The hardware of these four machines are varied. One machine has 32 (logical) cores with 256GB RAM. The other three machines have 20 (logical) cores and 188GB RAM. All machines run on Ubuntu 12.04LTS system.

### Reference genome analysis

We generated 31-mer sets for GRCh37 and GRCh38 by extracting 31-mers starting on every position (forward strand) in both assemblies for all autosomes and gonosomes. Any 31-mers containing IUPAC ambiguity codes were discarded. The 31-mers were queried against the population BWT to check for support in the 1000GP read set (forward and reverse orientation). The population BWT 31-mers (used in Figure 2a and Figure 2b) were generated from the final corrected set of read sequences.

### 1000 Genomes variant 31-mers

For each individual in 1000GP, we created a maternal and paternal genome by substituting the phased variants and generated 31-mers that overlap with every non-reference position. We excluded unphased, non-diploid (except gonosomal hemizygous), or conflicting variants (e.g. SNPs in regions which are also called as being deleted on the same chromosome copy), variants for which the exact coordinates could not be determined, reference alleles where an individual chromosome copy was contradictory (e.g. a region genotyped as reference for a deletion that also contains another non-reference variant) and filtered reference alleles that collided with each other by discarding all downstream reference loci within the overlapping region. In total, 0.16% of the variants were excluded. For each of the resulting haploblocks every contained 31-mer was generated and queried against the Population BWT.

### SNP genotyping

We used the 1000GP Illumina Omni chip data produced at the Broad institute (ftp://ftp.1000genomes.ebi.ac.uk/vol1/ftp/technical/working/20131122_broad_omni/Omni25_genotypes_2141_samples.b37.v2.vcf.gz) for the list of gold standard SNP genotypes. There were 1668 samples in the Omni chip genotypes that were included in the phase 3 1000GP freeze. Genotyping was carried out with the population BWT, by generating a reference and alternate allele using 99bp of flanking sequence for each site. We tiled each allele sequence with 34-mers with a step of 10bp. We queried the population BWT with the 34-mers and carried out a local Smith-Waterman alignment (match +1, mismatch penalty −4, gap open penalty −6, gap extension penalty −1) of the returned reads onto the reference and alternate allele, excluding divergent hits (if only mismatches, then allow maximum of three mismatches, otherwise allow a maximum of one indel and eight points penalty). Using the number of reads supporting the reference or alternative alleles, we assigned a genotypes according to Table 1. Finally, we output a new VCF file with the population BWT determined genotypes.

**Table 1.**
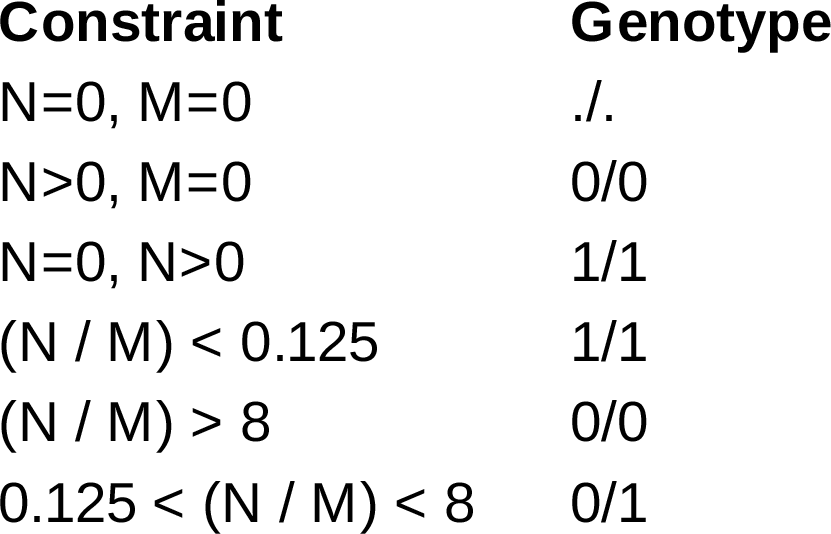
The population BWT SNP genotype assignment scheme. ‘N’ is number of reference supporting reads and ‘M’ number of alternative allele supporting reads, ‘.’ means unknown genotype, ‘0’ is the reference allele, and ‘1’ is the first alternative allele.

### Indel genotyping

We dowloaded a recent version of the Genome in a Bottle (GIAB) NA12878 variant set (v2.18,ftp://ftp-trace.ncbi.nlm.nih.gov/giab/ftp/release/NA12878_HG001/NISTv2.18/NISTIntegratedCalls_14datasets_131103_allcall_UGHapMerge_HetHomVarPASS_VQSRv2.18_all_nouncert_excludesimplerep_excludesegdups_excludedecoy_excludeRepSeqSTRs_noCNVs.vcf.gz) and filtered it to include only monoallelic indels. We then used Samtools/bcftools v1.1 (samtools mpileup-gut DP,DV,DP4,SP,DPR,INFO/DPR -EQ 0 -p -b [NA12878_BAM_FOFN] -f [GRCh37_REF] -l [NA12878_VARIANTS_BED] | bcftools call -mf GQ,GP -O z) and GATK v3.5 (java7 -jar-Xmx28G GenomeAnalysisTK.jar -R [GRCh37_REF] -I [NA12878_BAM_LIST] -L [NA12878_VARIANT_INTERVALS_BED] -T HaplotypeCaller -stand_call_conf 4--genotyping_mode GENOTYPE_GIVEN_ALLELES --output_mode EMIT_ALL_SITES --alleles [NA12878_VARIANTS_VCF]) (intervals are the indel start & end position padded by 150bp) to call and/or genotype those variants based on NA12878 low coverage and exome sequencing data. The Samtools/Bcftools calls were subsequently left normalized as we did not specify the alleles during the genotyping stage. The GATK results did not require this step. For population BWT genotyping, we generated a reference and alternate sequence for each indel by adding 100bp of flanking sequence to either the reference or alternate allele. We then generated 25-mers (1bp step) from these sequences and queried the population BWT for matching reads. Any 25-mer with greater than 100,000 matches or with a homopolymer length greater than 14bp were excluded. We further generated flanking sequences between 100-200bp upstream and downstream of each variant. If a 25-mer from these regions was found in any of the reads returned from the BWT we considered that as evidence that the corresponding read is either pointing away from the variant or too far away to overlap it and hence discarded the read. All remaining reads were collapsed into a non-redundant read set and aligned against both the reference and alternative alleles using exonerate v2.2.0 (--model ungapped --dnawordlen 25 --percent 90 --bestn 1). Alignment hits were subsequently filtered for reads that reach at least 2nt from the flank into the variant locus. Each valid read was then assigned to either reference or alternative allele based on the highest alignment score, with reads with equal scores being discarded. Finally, the full sample metadata was retrieved and the indels genotyped per sample using the read count thresholds in Table 1.

### Viral genome analysis

We downloaded 257,943 viral sequences from the CoreNucleotide division (http://www.ncbi.nlm.nih.gov/nuccore/on 20/02/2015) of GenBank (search string: "((((((txid10239[Organism]) AND 2000[SLEN]:300000[SLEN]))))) NOT patent"). We generated the virus taxon-specific 31-mers using Kraken v0.10.5-beta (Wood and Salzberg 2014) by generating a database containing fully assembled virus ("kraken-build --download-library viruses"), bacteria ("kraken-build --download-library bacteria"), and plasmid ("kraken-build --download-library plasmids"), and the GRCh38 human reference assembly ("kraken-build --download-library human") (built on 16/03/2015). We used this Kraken database to classify the virus sequences downloaded from GenBank. Of the 257,943 input sequences, 244,656 (94.8%) could be classified, 243,123 (94.3%) as viruses covering 4093 of 5808 virus taxa (70.5%). From the Kraken output files, we extracted 102,655,127 taxon-specific 31-mers. We queried the population BWT with these 31-mers, returning counts for the number of matching reads (query time 2d3h48'26" CPU time, 2d16h5'7" wall clock time using 80 threads). Of the 102.6M 31-mers, 435,799 from 886 taxa had matches in the Population BWT. 1,369 of the 31-mers match very large numbers of reads (>100,000), indicating that these contain little information and match repetitive or low complexity sequences, were discarded. We subsequently did full read sequence retrieval queries for the remaining 434,430 31-mers (0d5h2'19" CPU time, 0d5h9'22" wall clock time, using 10 threads). All reads returned from the population BWT were collapsed into a non-redundant set of sequences per taxon ID resulting in a final size of 113,193,726 reads.

Although we can be sure that each read contains at least one taxon specific 31-mer, this could be due to one or more sequencing errors in the 31-mer. Therefore, we reclassified the reads by short read alignment to the genome sequences using Smalt v0.7.5.1 (http://www.sanger.ac.uk/science/tools/smalt-0), which enabled us to examine the relative alignment score of matches to assess the classification. Based on the alignment results we chose a threshold of 75% of the maximum alignment score per read and included only reads that exceeded this threshold when aligning to a virus genome while staying below for any other kind of target sequence (human, bacteria, or plasmid). Each read fulfilling these criteria was then assigned to the virus it aligned best to. In case of equal best matches to different virus genomes one was chosen at random. Using this filter, 107,234,569 reads (94.7%) could be assigned to a virus covering 289 virus taxa. To assign samples, we queried the population BWT metadata database for sample information per read (total run time was 7d17h25'32" CPU time, 8d0h42'13" wall clock time).

For the samples found to contain HTLV-1, we downloaded the original fastq files from the 1000GP ftp site (http://ftp://ftp.1000genomes.ebi.ac.uk/vol1/http/phase3/data/) and aligned all of the reads using bwa mem v0.7.12 to a reference genome containing GRCh38 + HTLV-1. Table 2 gives the relative read counts for reads found to contain HTLV-1 from the BWT queries and alignment of the reads (no minimum mapping quality or length threshold for hits).

**Table 2.** The population and sample identifiers for the individuals found to contain evidence for HTLV-1 viral integrations. The read count columns indicate the number of viral reads by searching the population BWT and by re-alignment of all of the sequencing reads from the individuals to a GRCh38 * HTLV-1 reference genome.

| Population | Sample | Centre | Read counts |  |  |
| --- | --- | --- | --- | --- | --- |
|  |  |  | BWT | Alignment | Unique integrations |
| ACB | HG02325 | Broad institute | 1 | 2 | n/a |
| CLM | HG01357 | Illumina, BCM | 224 | 282 | 8 |
| ESN | HG03370 | WUGSC, WTSI | 40 | 92 | 2 |
| GWD | HG02675 | Broad institute | 104 | 123 | 3 |
| PEL | HG01918 | BGI | 19 | 32 | n/a |
| PEL | HG01917 | BGI | 21 | 34 | 2 |

## Data Access

All of the sequencing data used in this study is available from the 1000GP site under the accessions given in ftp://ftp.1000genomes.ebi.ac.uk/vol1/ftp/phase3/20130502.phase3.sequence.index. The collection of software used to build the population BWT server is available on Github (https://github.com/wtsi-svi/ReadServer).

## Acknowledgments

This work was supported by the Wellcome Trust. We thank John Marshall (Wellcome Trust Sanger Institute) for providing technical help and support for several aspects of this project.

## Author contributions

JTS, SMC, RD, ZI and TMK initiated the ideas for the study. DDD, ZL, SMC, and TK carried out the software development and computational analysis. MC provided advice for the virus genome analysis. DDD and TMK drafted the manuscript. All authors read and approved the final manuscript.

---

**Supplementary Figure 1:** Server diagram where each grey box represents a physical machine. The web server application that handles k-mer requests is Mongoose (https://github.com/cesanta/mongoose) and communication within the server is via ZeroMQ (https://github.com/zeromq). Users send k-mers and the type of output (matching read counts, matching read sequences, or matching reads with sample metadata). The output is returned in JSON format.

**Supplementary Figure 2:** Analysis of not supported regions in GRCh37. (a) Size distribution of regions covered by unsupported reference 31-mers (light green: exclusive to GRCh37, dark green: shared with GRCh38) (b) Bar plot showing the fraction of unsupported regions in the GRCh37 reference assembly containing high or low allele frequency variants. Significance testing was based on selecting 1,000 random regions from the genome.

**Supplementary Figures 3-8:** Viral sequences found in the 1000GP population BWT, one figure per continent. Individuals are represented on the x-axis. Only virus taxa with greater than 10 reads (black bars) in at least one sample are shown. Samples are ordered by population, source (blood: pink, LCL: dark green, unknown: light green), and total virus load (sum of read counts per virus and sample).

**Supplementary Table 1:** The 16 population BWT partitions, based on sequence suffix. Total number of sequences, length, BWT size, index size, and memory usage are given.

**Supplementary Table 2:** 1000GP population BWT server performance comparison for different values of k returning either matching read counts, matching read sequences, or read meta-data running on the internal network at the Wellcome Trust Sanger Institute.

**Supplementary Table 3:** Results for 1000GP population BWT SNP and indel genotyping for different size k-mers for all chromosome 20 sites compared to the 1000GP Illumina Omni chip and Genome in a Bottle (indels, NA12878), respectively.

**Supplementary Table 4:** Genes overlapped by not supported 31-mers in GRCh37 and/or GRCh38. Only genes were selected for which a region annotated as CDS in GENCODE 19 (for GRCh37) and/or GENCODE 23 (for GRCh38) was hit by a 31-mer.

